# ELLA-mix: Detecting Cell-to-Cell Heterogeneity in Subcellular RNA Localization

**DOI:** 10.64898/2026.11.23.768697

**Authors:** Aravind Bandreddi, Scott A. Bruce, Jade Wang

## Abstract

Subcellular RNA localization can differ between cells expressing the same gene, yet most analyses summarize transcript positions with a single gene-level pattern. We propose ELLA-mix, a finite mixture model for identifying cell subpopulations with distinct radial localization patterns from imaging-based spatial transcriptomics data. Transcript positions are normalized relative to the cell boundary and modeled using flexible spline densities within a count-conditioned nonhomogeneous Poisson process framework. Conditioning on each cell’s transcript count separates differences in localization from differences in abundance. We estimate component densities and cell memberships using an expectation-maximization algorithm and select the number of patterns using the Bayesian information criterion, restricting consideration to models with stable cell assignments and sufficiently distinct components. Simulation studies evaluated the type I error and power of the test for nonuniform localization, as well as the accuracy of selecting the number of mixture components and estimating their radial distributions. ELLA-mix correctly selected the number of components in most simulated settings. In seqFISH+ mouse fibroblasts, ELLA-mix identified distinct cell subpopulations for genes previously assigned a single localization class; genes showing such variation were enriched for endoplasmic reticulum and extracellular matrix functions. In Xenium ovarian cancer tissue, more peripheral localization patterns were associated with higher proportions of proliferating cells, particularly in S phase. These results show how modeling cell-to-cell variation can reveal localization patterns and biological associations obscured by gene-level summaries.

## 1. Introduction

Spatial transcriptomics encompasses a rapidly growing collection of technologies that measure gene expression while preserving spatial context in tissues. Recent technological advances have enabled spatial transcriptomic measurements at subcellular resolution. Imaging-based platforms, including MERFISH, seqFISH+, Xenium, and CosMx, map individual RNA molecules in situ (Chen et al., 2015; Eng et al., 2019; 10x Genomics, 2026b; NanoString Technologies, 2026), whereas sequencing-based platforms such as Stereo-seq and Visium HD quantify gene expression in fine spatial bins (Chen et al., 2022; 10x Genomics, 2026a). These advances have made it possible to study not only single cell transcript abundance but also the spatial distribution of RNA molecules within individual cells. Subcellular mRNA localization is an important mechanism of spatial gene regulation: positioning transcripts within a cell can affect where proteins are synthesized and how the cell responds to local cues (Das et al., 2021). For example, *β*-actin mRNA localizes near the leading edge of migrating fibroblasts, and disrupting this localization impairs cell polarity and directional movement (Shestakova et al., 2001). Such observations motivate the analysis of subcellular distributions, such as enrichment near the nucleus or cell membrane, that cannot be inferred from total expression alone.

Despite the biological importance of subcellular mRNA localization, relatively few methods and tools have been developed to characterize these patterns from spatial transcrip-tomics data. Bento assigns predefined localization labels to gene-level transcript distri-butions, while SPRAWL uses rank-based statistics to quantify peripheral and central localization (Mah et al., 2024; Bierman et al., 2024). HiVis analyzes subcellular localization in Visium HD data using image-derived compartment annotations (Novoselsky et al., 2026). FISHFactor identifies latent subcellular expression patterns across genes and cells, while SubCellSpace represents gene-specific localization patterns in individual cells and examines their variation across cells (Walter et al., 2023; Wouters et al., 2026). SVC uses a vision transformer to learn cell-specific subcellular transcript organization and spatial dependencies across genes (Wan et al., 2026). ELLA models the radial distribution of a gene’s transcripts and tests for subcellular spatial variation, estimating a shared localization pattern across the analyzed cells (Wang and Zhou, 2025).

Moreover, subcellular localization may vary among cells expressing the same gene, even within an annotated cell type. SubCellSpace characterizes gene-specific localization patterns and their variation across individual cells, but its gene-level test assesses nonrandom spatial localization rather than the presence of distinct localization patterns across cells (Wouters et al., 2026). In seqFISH+ fibroblasts, ELLA revealed cell-cycle-associated differences in radial localization by fitting the model separately to cells assigned to the G1, S, and G2M phases (Wang and Zhou, 2025). However, this analysis required cell groups to be defined in advance. When the relevant groups are unknown, pooling cells may obscure distinct localization patterns, whereas estimating a separate pattern for each cell can be unreliable because individual cells often contain few observed transcripts. Furthermore, detecting nonuniform localization does not establish whether a gene exhibits a single shared pattern or multiple distinct patterns across cells. These considerations motivate a mixture-model framework for characterizing cell-to-cell heterogeneity in radial localization and selecting the number of distinct localization patterns.

To capture variation across cells while estimating patterns shared among them, we introduce a mixture model for the radial locations of a gene’s transcripts. Specifically, we use a count-conditioned nonhomogeneous Poisson process (NHPP) mixture, with flexible spline densities to characterize component-specific radial localization patterns. The model estimates the component densities and mixing proportions, together with posterior probabilities of component membership for individual cells. By conditioning on the observed transcript counts, the model accommodates overdispersion in transcript abundance across cells while focusing inference on spatial localization. We estimate the model parameters by maximum likelihood using an expectation-maximization (EM) algorithm. We use a likelihood-ratio test to assess the null hypothesis of spatial uniformity against a nonuniform alternative. For genes exhibiting nonuniform localization, we select the number of mixture components using the Bayesian information criterion (BIC), restricting consideration to models with stable cell assignments and sufficiently distinct component densities.

The remainder of the paper is organized as follows. Section 2 presents the conditional mixture model for radial transcript locations. Section 3 describes model fitting and inference for the number of localization patterns. Section 4 evaluates the method through simulations, and Section 5 applies it to spatial transcriptomics data. Section 6 concludes with a discussion.

## 2. Model Formulation

### 2.1 Data structure

We analyze one gene at a time and suppress the gene index throughout. Suppose the gene is observed in *i* = 1, …, *n* cells, and cell *i* contains *m*_*i*_ observed transcripts (*m*_*i*_ ⩾ 1), indexed by *j* = 1, …, *m*_*i*_. For each cell, the observed spatial information consists of its segmented region *C*_*i*_ with boundary *∂C*_*i*_, its nucleus center **c**_*i*_ = (*c*_*i*1_, *c*_*i*2_)^*⊤*^, computed as the mean of the vertices of the segmented nucleus boundary, and the transcript coordinates **x**_*ij*_ = (*x*_*ij*1_, *x*_*ij*2_)^*⊤*^ *∈ C*_*i*_.

Let *r*_*ij*_ = ∥**x**_*ij*_ *−* **c**_*i*_∥_2_ be the distance from transcript *j* to the nucleus center in cell *i*, and let *R*_*ij*_ be the distance from **c**_*i*_ to the first intersection with *∂C*_*i*_ of the ray from **c**_*i*_ through **x**_*ij*_. The normalized radial location is

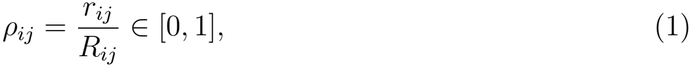

where *ρ*_*ij*_ = 0 corresponds to the nucleus center and *ρ*_*ij*_ = 1 to the cell boundary. Transcripts with *ρ*_*ij*_ *>* 1, which lie outside the segmented cell boundary, are excluded. This normalization places transcripts from cells of different sizes and shapes on a common radial scale. Thus, the data supplied to our model are the normalized radial locations *D*_*ρ*_ = *{ρ*_*ij*_ : *j* = 1, …, *m*_*i*_, *i* = 1, …, *n}*.

### 2.2 Conditional mixture model for radial localization patterns

Our objective is to characterize cell-to-cell heterogeneity in subcellular RNA localization by identifying distinct radial localization patterns and estimating their prevalence across cells. We model transcript locations conditional on each cell’s observed transcript count *m*_*i*_. For a specified number *K* of mixture components, let *z*_*i*_ *∈ {*1, …, *K}* denote the latent component membership of cell *i*. Given *z*_*i*_ = *k*, we model the normalized radii 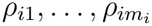 of cell *i* as an NHPP on [0, 1], with the mean measure over an interval [*a, b*] given by

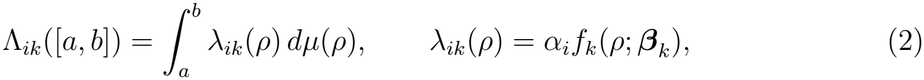

where 0 ⩽ *a* ⩽ *b* ⩽ 1, and

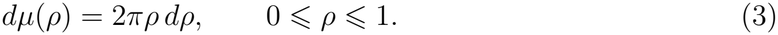

Here, *α*_*i*_ *>* 0 is a cell-specific scale parameter, and *dµ*(*ρ*) = 2*πρ dρ* is the area of the thin ring at radius *ρ* in the normalized unit disk. Defining *λ*_*ik*_ with respect to this measure makes it an intensity per unit area, so a flat *f*_*k*_ corresponds to transcripts spread uniformly over the cell. The intensity *λ*_*ik*_ and density *f*_*k*_ are defined with respect to this measure. W impose the normalization constraint 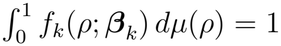, so that *f*_*k*_ integrates to 1 over the area of the normalized cell and is a probability density of transcript locations. Hence *f*_*k*_ describes only the shape of the localization pattern, while Λ_*ik*_([0, 1]) = *α*_*i*_ is the expected number of transcripts in cell *i*.

Conditional on *m*_*i*_ and *z*_*i*_ = *k*, the transcript radii are independently distributed with density, with respect to *µ*,

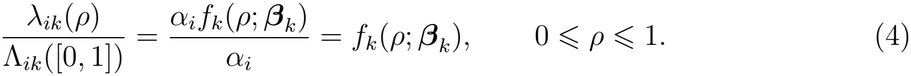

Thus, the cell-specific scale parameter *α*_*i*_ is eliminated by conditioning on the observed transcript count.

Let **B**(*ρ*) = (*B*_1_(*ρ*), …, *B*_*P*_ (*ρ*))^*⊤*^ denote a vector of *P* cubic B-spline basis functions evaluated at the normalized radial position *ρ ∈* [0, 1], and let ***β***_*k*_ *∈* ℝ^*P*^ denote the corresponding coefficient vector for component *k*. In all analyses we use *P* = 5, with a single interior knot at *ρ* = 0.5. We model the normalized radial density as

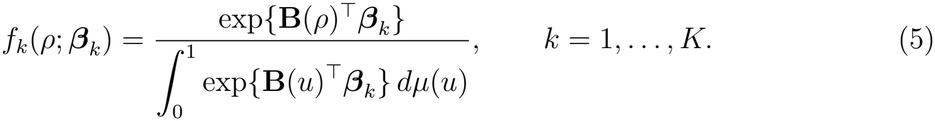

The normalized density is invariant to a common shift in the spline coefficients, so ***β***_*k*_ is identifiable only up to an additive constant.

Let ***π*** = (*π*_1_, …, *π*_*K*_)^*⊤*^ denote the mixing proportions, where *π*_*k*_ *>* 0 and 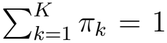. We specify component membership conditional on the observed counts as

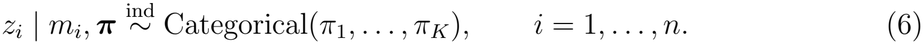

The mixing proportion *π*_*k*_ represents the proportion of cells belonging to component *k*. All transcripts within a cell share the same *z*_*i*_, making the cell the unit of clustering. Thus, *K* = 1 represents one radial pattern shared across cells, whereas *K >* 1 allows distinct localization patterns across cell subpopulations.

Let **Θ**_*K*_ = (***π, β***_1_, …, ***β***_*K*_) denote the collection of unknown parameters in the conditional mixture model. Summing over the latent component memberships, the likelihood of the observed radial transcript locations, conditional on the cell-level transcript counts, is

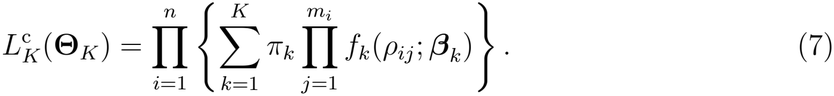

The model assumes that, conditional on *m*_*i*_ and *z*_*i*_, transcript locations within each cell are independent draws from the corresponding component density.

## 3. Estimation and Inference

### 3.1 Estimation for a fixed number of patterns

For a specified number *K* of mixture components, we estimate the model parameters by maximizing the conditional likelihood in (7) using EM algorithm. We use *P* cubic B-spline basis functions to parameterize the component densities, with the same basis across components and candidate values of *K*. Let *Z*(***β***_*k*_) denote the normalizing constant in (5) and **s**_*i*_ the cell-level sufficient statistic,

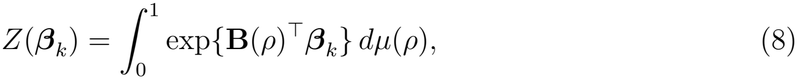

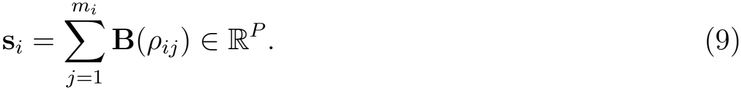

The data from cell *i* enter the likelihood only through **s**_*i*_ and *m*_*i*_. Substituting (5) into the inner product of (7), the contribution of cell *i* under component *k* is

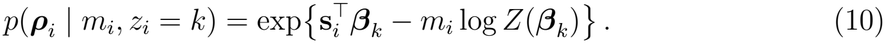

Marginalizing over the latent component memberships yields the conditional observed-data log-likelihood

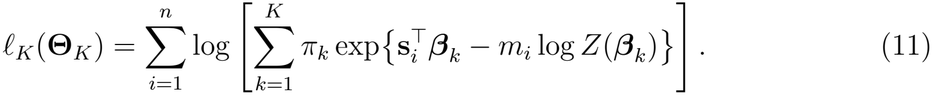

Let *w*_*ik*_ = *I*(*z*_*i*_ = *k*) denote the indicator that cell *i* belongs to component *k*. If the memberships were observed, combining (10) with (6) would give the conditional completedata log-likelihood

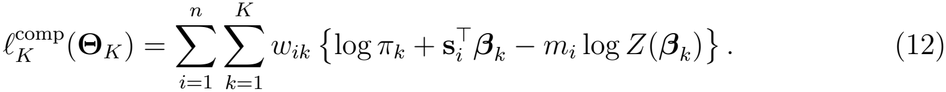

In the E-step, we compute the posterior probability that cell *i* belongs to component *k*, conditional on its observed transcript locations and count, as

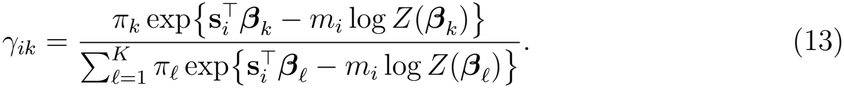

In the M-step, the mixing proportions are updated in closed form as

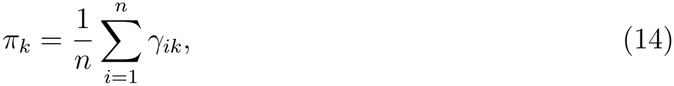

and the spline coefficient vector ***β***_*k*_ for each component *k* is updated by maximizing

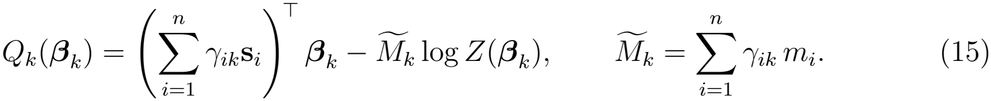

Here, 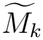 denotes the responsibility-weighted total number of transcripts assigned to component *k*. The objective function *Q*_*k*_(***β***_*k*_) is smooth and concave, with gradient given by

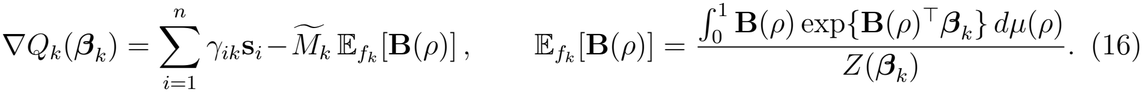

We maximize *Q*_*k*_ using L-BFGS with the analytic gradient in (16). The integrals in (8) and (16) are approximated numerically on a fixed grid of 200 radial points, with area weights 2*πρ* Δ*ρ*. The EM algorithm is initialized with equal mixing proportions and small random spline coefficients. Iterations terminate when the relative change in the observed-data log-likelihood falls below 10^*−*6^ or the maximum of 500 iterations is reached. After fitting, each cell is assigned to the component with the highest posterior membership probability, 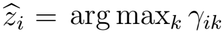.

### 3.2 Testing for nonuniform localization

We test for nonuniform radial localization against complete spatial randomness (CSR) using a likelihood-ratio test. Under CSR, the intensity is constant on the unit disk, *λ*(*r*) = 1*/π*, yielding a conditional log likelihood of *l*_0_ = *−N* log *π*, where *N* is the total number of transcripts for a gene. Since CSR is nested within the single-pattern model (*K* = 1), the likelihood-ratio statistic is

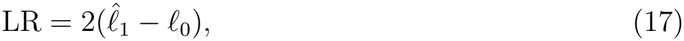

where 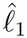 is the maximized log likelihood under *K* = 1. Under CSR, LR is asymptotically distributed as 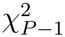, where *P −*1 is the number of free spline parameters. Simulations confirm good calibration under CSR (Figure 1a). We apply the Benjamini-Hochberg procedure to control the false discovery rate at 5% across genes. Genes with significant nonuniform localization are subsequently considered for mixture-component selection.

**Figure 1.**
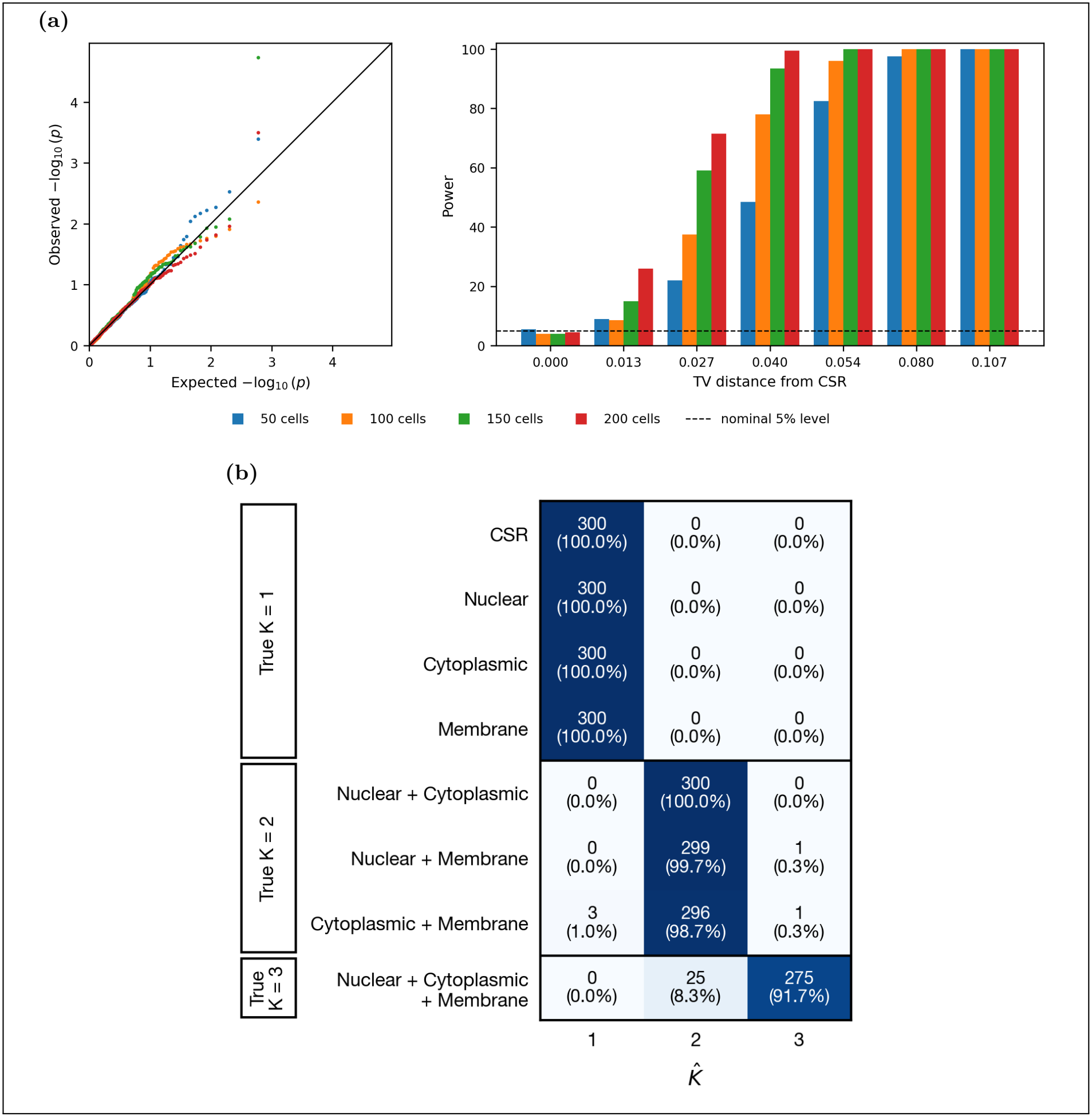
(a) Left: QQ plot of the *p*-values of the likelihood-ratio test for nonuniform localization under CSR, with 50, 100, 150 and 200 cells per gene (300 simulated genes per cell count). Right: empirical power of the test across TV distances from CSR for the membrane localization pattern (200 simulated genes per bar). Bars at TV = 0 are CSR genes and give the type I error; the dashed line marks the nominal 5% level. (b) Selected number of localization patterns 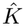across the 8 simulation scenarios (300 genes per scenario, 150 cells per gene). Each row gives the number and percentage of genes assigned 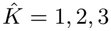; dark cells on the diagonal are correct selections.

### 3.3 Selection of the number of localization patterns

We select the number of mixture components using the Bayesian information criterion (BIC). For each gene, we fit the conditional mixture model with *k* = 1, …, *K* components, where *K* denotes the maximum number of components considered. For each candidate *k*, the model is fitted from three random initializations, and the fit with the highest observed-data log-likelihood, denoted by 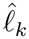, is retained. Multiple initializations reduce sensitivity to local maxima and allow us to assess the stability of the estimated cell assignments.

To exclude models with unstable cell assignments or poorly separated component densities, we define an admissibility indicator *A*_*k*_ *∈ {*0, 1*}*. A model with *k* ⩾ 2 is considered admissible (*A*_*k*_ = 1) if its cell assignments are sufficiently stable across initializations and its fitted component densities are sufficiently distinct. Otherwise, *A*_*k*_ = 0. The single-component model is always admissible (*A*_1_ = 1). Details of the admissibility criteria are provided in Supplementary Section S1.

Among the admissible models, we select the number of components by minimizing

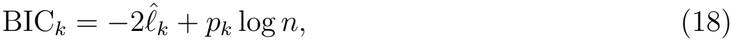

where *n* denotes the number of cells and *p*_*k*_ is the number of identifiable model parameters. Each component contributes *P −* 1 free spline coefficients, since adding a common constant to the coefficients leaves the normalized density unchanged. Together with the *k −* 1 free mixing proportions, the total number of parameters is *p*_*k*_ = *k*(*P −* 1) + (*k −* 1) = *kP −* 1. The selected number of components is therefore

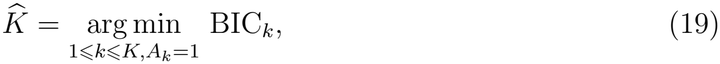

with ties resolved in favor of the smaller model.

## 4. Simulations

We conducted simulation studies to evaluate the type I error and power of ELLA-mix in detecting nonuniform RNA localization, as well as its accuracy in selecting the number of localization patterns and estimating their radial distributions. We considered three representative radial localization patterns, nuclear, cytoplasmic and membrane, ranging from enrichment near the nucleus to enrichment near the cell boundary. Details of the simulation design and data generation are provided in Supplementary Section S2.

Under the null hypothesis of complete spatial randomness (CSR), we generated transcript locations uniformly over the unit disk to assess the calibration of the likelihood-ratio test for nonuniform localization. The test compares the uniform model with a flexible single-component spline model, using a *χ*^2^ reference distribution with *P −* 1 degrees of freedom. We evaluated the distribution of the resulting *p*-values and the empirical type I error at the nominal significance level of 0.05, using 300 CSR genes for each of 50, 100, 150 and 200 cells per gene. The empirical type I error was 5.7%, 8.7%, 6.3% and 4.3%, respectively, and the *p*-values were well calibrated (Figure 1a).

Under the alternative hypothesis, we evaluated the power of ELLA-mix to detect nonuni-form localization. We generated transcript distributions with varying pattern strengths, quantified by their TV distances from CSR, and varied the number of cells per gene. Figure 1a shows how power varies with pattern strength for 50, 100, 150 and 200 cells per gene, using the membrane localization pattern as an example. Power increased with both pattern strength and the number of cells. For example, at a TV distance of 0.040, power was 49%, 78%, 94% and 100% with 50, 100, 150 and 200 cells, and all genes rejected uniformity at TV distances of 0.107 or more. Similar trends were observed for the other localization patterns.

Next, we evaluated the accuracy of ELLA-mix in selecting the number of localization patterns. We considered 8 simulation scenarios with one, two, or three underlying patterns, with all cells sharing a pattern following the same radial distribution and 150 cells per gene. Figure 1b summarizes the selection results. Among the 1,200 genes generated with a single pattern, including the 300 CSR genes, ELLA-mix correctly selected one component for all 1,200 (100%) genes. For the 900 genes generated with two patterns, ELLA-mix correctly selected two components for 895 (99.4%) genes; of the remaining 5, 3 were assigned one component and 2 were assigned three. Among the 300 genes generated with three patterns, 275 (91.7%) were correctly assigned three components, and the remaining 25 (8.3%) were assigned two. These results demonstrate that ELLA-mix accurately selected the number of localization patterns in all scenarios.

Lastly, we evaluated the accuracy of ELLA-mix in estimating the radial localization patterns. For genes with the correctly selected number of components, we matched the estimated components to the true patterns and measured the estimation error using the total variation (TV) distance between their normalized radial densities. With 150 cells per gene, the mean TV distances were 0.018, 0.029, and 0.039 for genes with one, two, and three components, respectively (Table 1). The estimation error decreased steadily with the number of cells, from 0.028 to 0.016 for single-pattern genes between 50 and 200 cells. Errors were larger for genes with more components, because each component is estimated from only a subset of the cells. With 100 or more cells per gene, the estimated patterns differed from the true patterns by less than 5% of transcripts on average in every scenario.

**Table 1.** Estimation accuracy of the fitted radial patterns. For simulated genes with the correctly selected number of components, each fitted component was matched to its closest true pattern, and the error was measured by the total variation (TV) distance between the fitted and true radial densities. Entries are the mean (SD) of the error across genes.

|  | Cells per gene |  |  |  |
|---|---|---|---|---|
| Scenario | 50 | 100 | 150 | 200 |
| CSR | 0.027 (0.011) | 0.019 (0.008) | 0.016 (0.006) | 0.014 (0.005) |
| Nuclear | 0.032 (0.011) | 0.024 (0.007) | 0.022 (0.005) | 0.020 (0.004) |
| Cytoplasmic | 0.027 (0.011) | 0.018 (0.008) | 0.015 (0.006) | 0.014 (0.005) |
| Membrane | 0.026 (0.010) | 0.021 (0.007) | 0.018 (0.005) | 0.016 (0.004) |
| Nuclear + Cytoplasmic | 0.052 (0.021) | 0.035 (0.010) | 0.030 (0.008) | 0.026 (0.007) |
| Nuclear + Membrane | 0.045 (0.013) | 0.032 (0.008) | 0.027 (0.006) | 0.025 (0.005) |
| Cytoplasmic + Membrane | 0.061 (0.029) | 0.039 (0.016) | 0.031 (0.011) | 0.027 (0.010) |
| Nuclear + Cytoplasmic + Membrane | 0.073 (0.024) | 0.048 (0.016) | 0.039 (0.010) | 0.034 (0.009) |

## 5. Data Analysis

We applied the proposed method to two spatial transcriptomics datasets representing distinct biological settings: seqFISH+ data from cultured mouse fibroblasts and Xenium Prime data from human ovarian cancer tissue. The fibroblast dataset allows us to investigate localization heterogeneity within a relatively homogeneous cell population, whereas the ovarian cancer dataset provides a more complex tissue setting in which localization patterns may vary across cell types and cellular states. For each gene, we retained cells with at least 10 observed transcripts and required at least 20 such cells for analysis.

### 5.1 seqFISH+

#### Data and gene filtering

We applied the method to a seqFISH+ dataset (Eng et al., 2019), which profiles transcripts at subcellular resolution in cultured NIH/3T3 mouse fibrob-lasts. The data consist of 179 cells imaged in 17 fields of view (Figure 2a), with 4.68 million transcripts from 7,095 detected genes. We used the cell and nucleus segmentations provided by Bento (Mah et al., 2024). Using these segmentations, we computed the normalized radial position *r ∈* [0, 1] for each transcript, where *r* = 0 corresponds to the nucleus centroid and *r* = 1 to the cell boundary.

**Figure 2.**
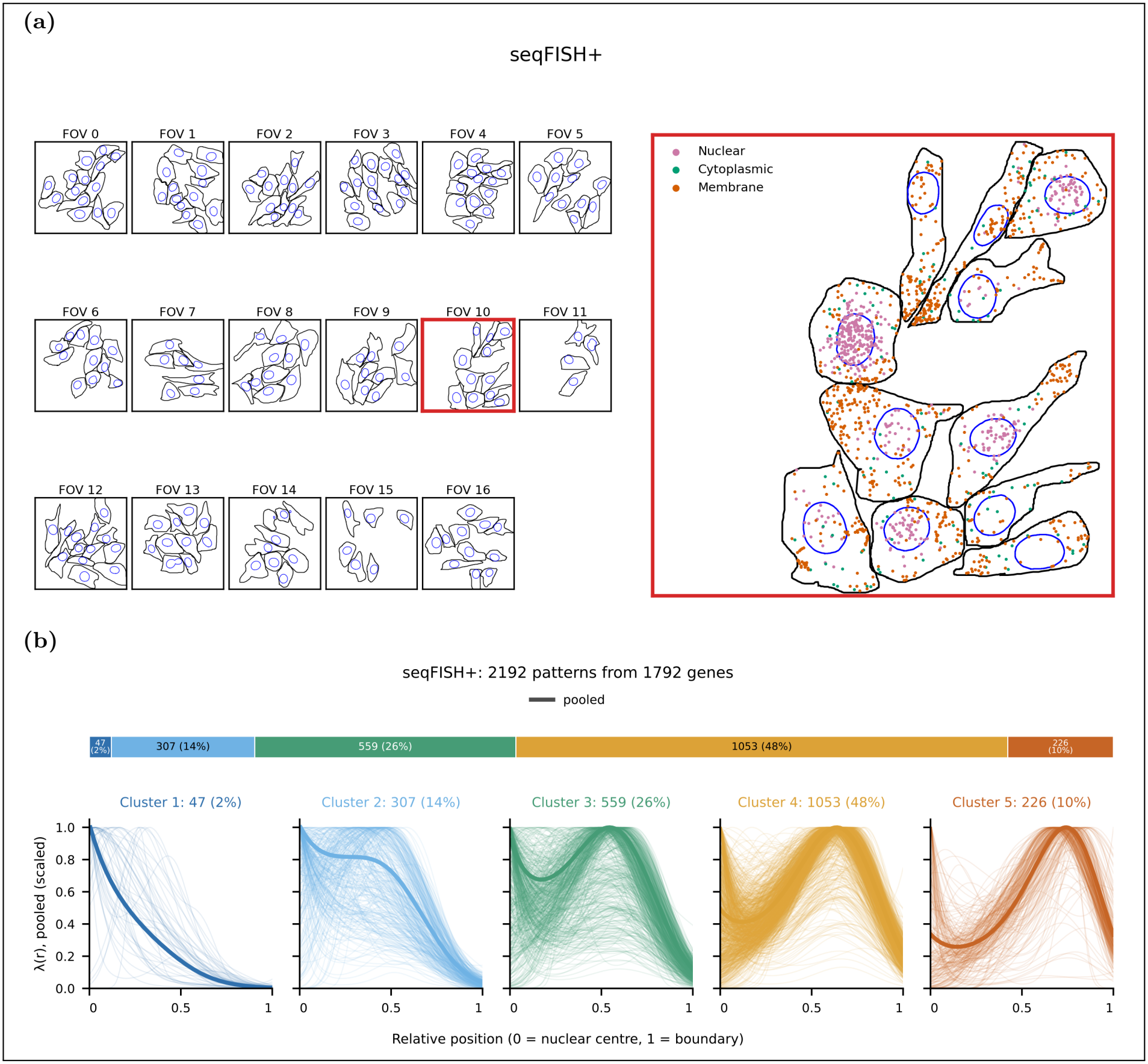
(a) The seqFISH+ dataset. Left: the 17 fields of view (FOVs), shown at the same scale, with cell boundaries (black) and nuclei (blue); the red box marks FOV 10. Right: FOV 10 enlarged, showing transcripts of three genes with nuclear, cytoplasmic and membrane patterns. (b) Estimated radial localization patterns in the seqFISH+ dataset. The fitted components of all genes are grouped into five pattern types by mean radial position (top bar: number and percentage of components per type). In each panel, thin lines are the intensities *λ*(*r*) of individual components and the thick line is the pooled pattern of the type, estimated from the transcripts of all cells assigned to its components with each component weighted equally. Curves are scaled to a maximum of 1.

#### Estimated localization patterns

We included 1795 genes in the analysis, of which 1792 showed significant nonuniform localization. Across the 1792 genes with localized patterns, ELLA-mix identified 2192 mixture components, each representing a radial localization pattern for a subpopulation of cells. To characterize the overall distribution of localization patterns, we classified the fitted components into five pattern categories according to their mean radial positions, ranging from the most nuclear (Pattern 1) to the most peripheral (Pattern 5). For each category, we obtained an average radial intensity curve, weighting each component equally (Figure 2b). Of the fitted components, 74% were classified into the cytoplasmic categories (Patterns 3 and 4), 10% into the peripheral category (Pattern 5), and 16% into the nuclear or perinuclear categories (Patterns 1 and 2).

#### Genes previously assigned a single pattern contain distinct subpopulations

Eng et al. (2019) classified each gene into a single subcellular localization class. Figure 3a compares the original localization classes with the ELLA-mix results for the 69 genes included in our analysis. Among the 47 genes for which ELLA-mix identified a single pattern, the estimated patterns were largely consistent with the original classifications (Figure 3b): 11 of 14 protrusion genes had a peripheral pattern (one, Net1, did not reject uniformity), 15 of 19 cytoplasm genes a cytoplasmic pattern, all 10 nuclear/perinuclear genes a perinuclear or inner cytoplasmic pattern, and 3 of the 4 genes in their first nuclear/perinuclear subcluster a nuclear pattern.

**Figure 3.**
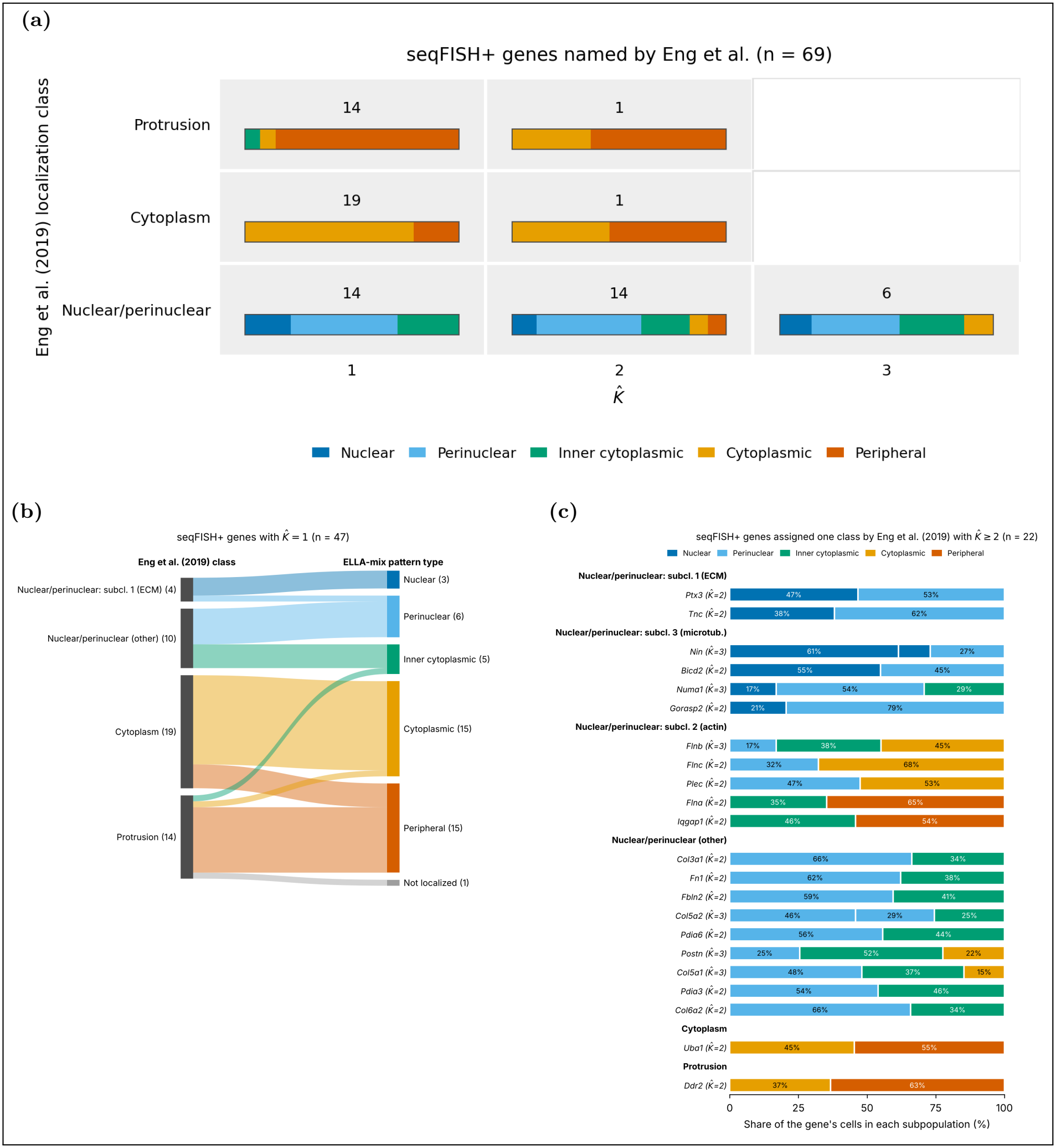
(a) Localization classes of Eng et al. (2019) compared with the selected number of patterns 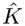 for the 69 seqFISH+ genes they classified that passed our filters; the nuclear/perinuclear class includes its three subclusters. Each cell gives the number of genes. The bar in each cell shows the pattern types of the fitted components of these genes, each component weighted by its share of cells 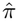, so that each gene contributes equally; because different genes can have different patterns, a cell can contain more pattern types than 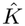. (b) Localization classes from Eng et al. (2019) (left) and ELLA-mix pattern types (right) for the 47 genes assigned a single pattern by ELLA-mix. Band widths are proportional to the number of genes. “Not localized” denotes a gene that did not reject uniformity. (c) ELLA-mix subpopulations for the 22 genes assigned two or three patterns. Each bar is one gene, grouped by its localization class in Eng et al. (2019); segments show the estimated proportion of cells in each subpopulation, coloured by its pattern type.

For the remaining 22 genes, ELLA-mix identified 2-3 distinct cell subpopulations, revealing localization heterogeneity not captured by the original gene-level classifications (Figure 3c).

For example, *Flnb*, originally classified as nuclear/perinuclear, was separated by ELLA-mix into three subpopulations across 169 cells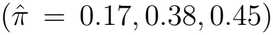 with nucleus, cyto-plasmic, and membrane patterns (Figure 4a; example cells in Figure 4b). Given the role of filamin B in actin cytoskeletal organization, these differences may reflect variation in cytoskeletal states among fibroblasts (Zhao et al., 2016). The other genes in the same subcluster showed similar heterogeneity: each of *Flna, Flnc, Iqgap1*, and *Plec* had one subpopulation closer to the nucleus and a larger one in the cytoplasm or near the cell boundary.

**Figure 4.**
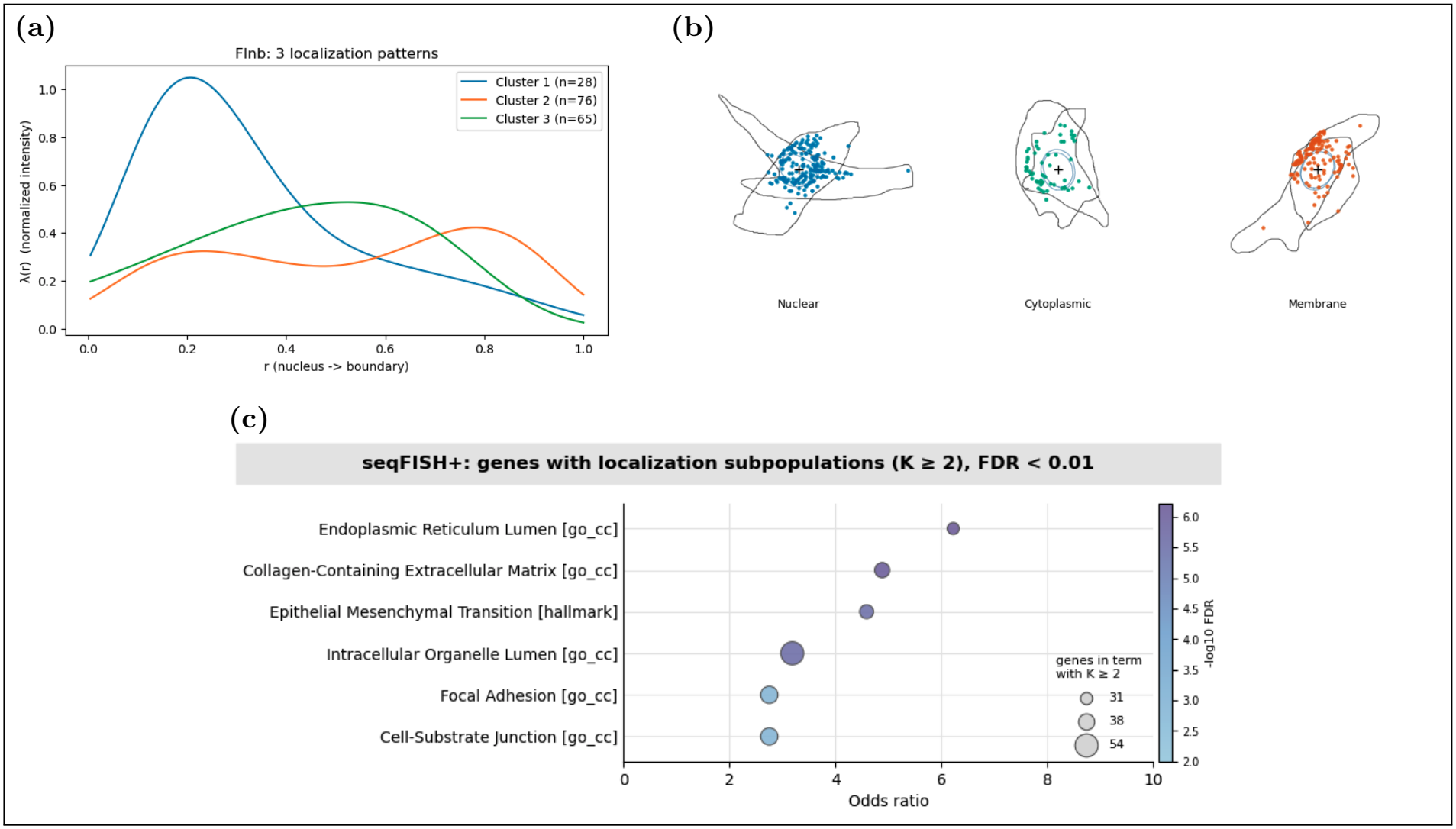
(a) Fitted radial localization patterns for *Flnb* (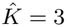; 169 cells), which Eng et al. (2019) classified as nuclear/perinuclear. (b) Example cells from each ELLA-mix subpopulation for *Flnb*. Each panel shows one cell with its boundary (black), nucleus (blue) and the gene’s transcripts (dots), coloured by subpopulation; the cross marks the cell centre. (c) Over-representation analysis of genes with 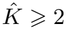against all 1,795 tested genes. Each dot is a gene set; its position gives the odds ratio, its size the number of genes with 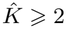 in the set, and its color the Benjamini–Hochberg *q*-value.

#### Over-representation analysis

To investigate the biological functions associated with cell-to-cell heterogeneity in radial localization, we performed over-representation analysis (ORA) of the 376 genes with multiple localization components 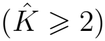, using Gene Ontology (GO) Cellular Component (CC) and Biological Process (BP) terms and MSigDB Hallmark gene sets, with all 1,795 tested genes as the background and Benjamini-Hochberg correction for multiple testing. Six gene sets were significantly enriched at an FDR below 0.01 (Figure 4c). The most strongly enriched sets included endoplasmic reticulum (ER) lumen (*p*_FDR_ = 6.0 *×* 10^*−*7^), collagen-containing extracellular matrix (*p*_FDR_ = 7.6 *×* 10^*−*7^), and Hallmark epithelial–mesenchymal transition (*p*_FDR_ = 2.9 *×* 10^*−*6^). The remaining enriched sets were intracellular organelle lumen, focal adhesion, and cell-substrate junction, which showed substantial overlap with the preceding sets or with one another. Genes contributing to these enrichments include ER chaperones (*Pdia3, Pdia6, Hyou1*) and collagens (*Col3a1, Col4a2, Col5a1, Col5a2, Col6a1, Col6a2*), which encode ER-resident proteins and secreted extracellular matrix proteins, respectively. These findings suggest that genes involved in the secretory pathway and cell-matrix adhesion are enriched among genes exhibiting cell-to-cell heterogeneity in radial mRNA localization. Such heterogeneity may reflect differences in the spatial organization of protein synthesis associated with secretion and cell-matrix interactions (Das et al., 2021).

### 5.2 Xenium

#### Data and gene filtering

We next applied ELLA-mix to a publicly available Xenium Prime 5K dataset of formalin-fixed paraffin-embedded (FFPE) human ovarian cancer tissue provided by 10x Genomics (10x Genomics, 2024). The tissue comprises diverse cell populations, including malignant, stromal, immune, and endothelial cells (Figure 5a). The dataset contains 407,124 cells and 106.5 million transcripts assigned to cells, from a panel of approximately 5,100 genes. The median number of transcripts per cell was 178 (IQR 66–388). Applying the gene-filtering criteria yielded 135 genes, detected in 22 to 128,581 cells.

**Figure 5.**
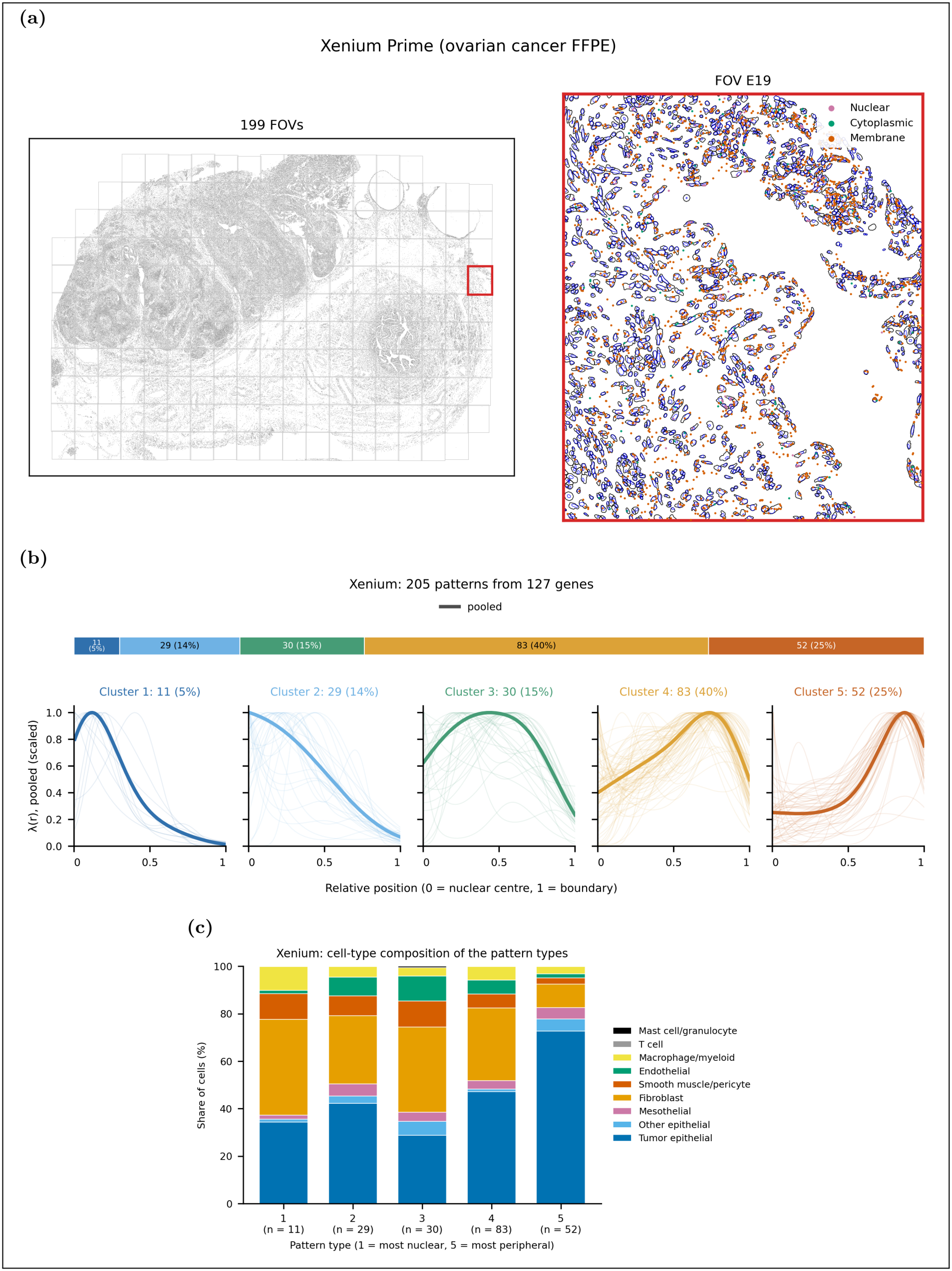
(a) The Xenium ovarian cancer dataset. Left: the tissue section and its 199 fields of view (FOVs); the red box marks FOV E19. Right: FOV E19 enlarged, with cell boundaries (black) and nuclei (blue), showing transcripts of three genes with nuclear, cytoplasmic and membrane patterns. (b) Estimated radial localization patterns in the Xenium dataset. The fitted components of all genes are grouped into five pattern types by mean radial position (top bar: number and percentage of components per type). In each panel, thin lines are the intensities *λ*(*r*) of individual components and the thick line is the pooled pattern of the type, estimated from the transcripts of all cells assigned to its components with each component weighted equally. Curves are scaled to a maximum of 1. (c) Cell-type composition of the five pattern types in the Xenium data. For each type, cells assigned to its mixture components were pooled across genes; bars show the percentage of these cells in each cell type. *n* is the number of components in each type (1 = most nuclear, 5 = most peripheral).

**Figure 6.**
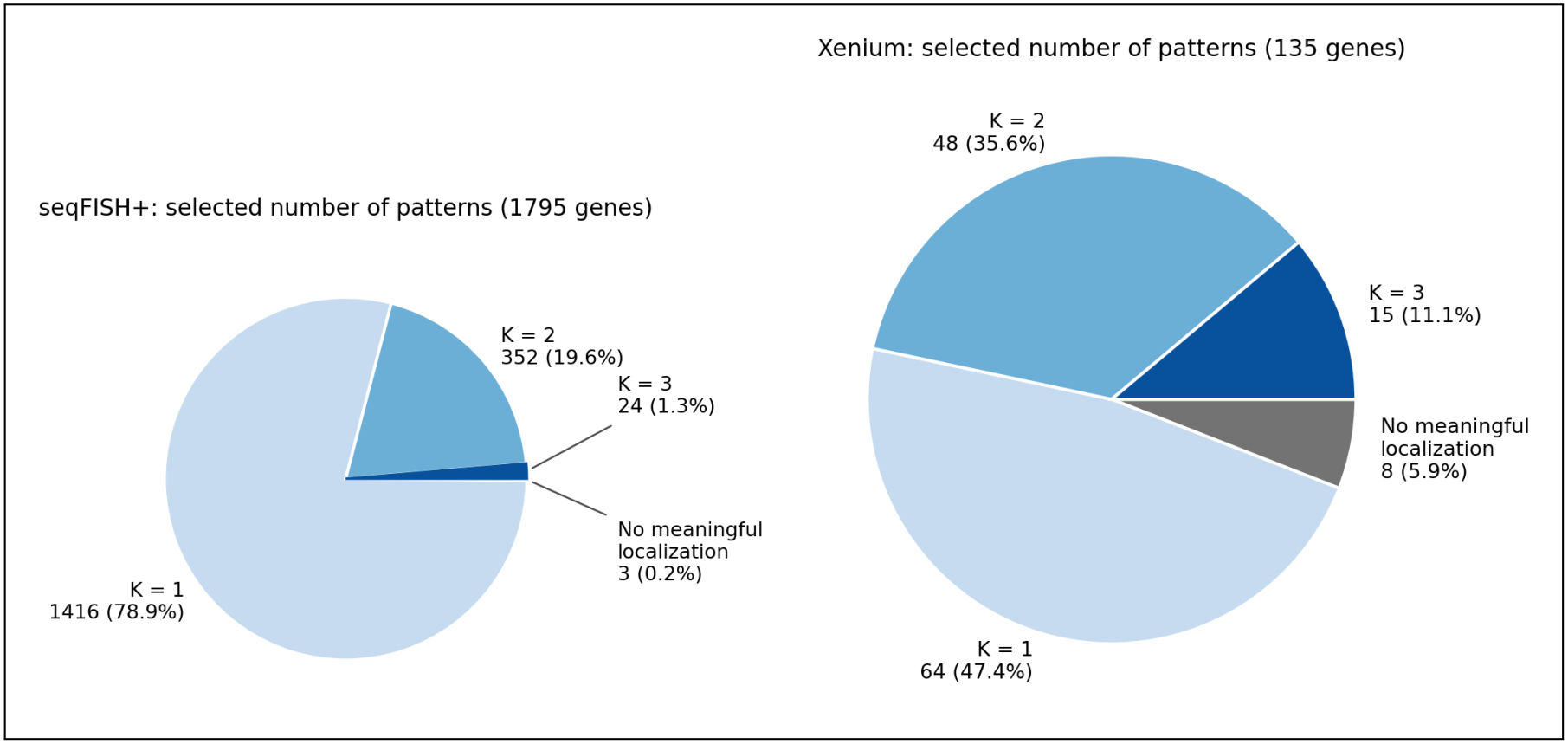
Selected number of localization patterns 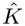 in the seqFISH+ (left; 1795 genes) and Xenium (right; 135 genes) datasets. Genes with 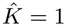 whose fitted pattern differed from a uniform spread by a total variation distance below 0.05 are shown as showing no meaningful localization.

#### Estimated localization patterns

We included 135 genes in the analysis, applying ELLA-mix to each gene across all eligible cells without stratifying by cell type. Of these genes, 127 (94.1%) showed significant nonuniform localization. Among these genes, ELLA-mix identified a single radial localization pattern for 64 (50.4%) genes, two patterns for 48 (37.8%), and three patterns for 15 (11.8%). Thus, 63 genes (49.6%) exhibited multiple localization patterns, compared with 21.0% in seqFISH+. This higher proportion may reflect the greater cellular diversity of ovarian cancer tissue. To characterize the distribution of localization patterns, we classified the 205 fitted mixture components into five pattern categories according to their mean radial positions, following the same procedure as in the seqFISH+ analysis. For each category, we obtained an average radial intensity curve, weighting each component equally (Figure 5b). Compared with seqFISH+, a higher proportion of components in Xenium were classified into the peripheral category (Pattern 5; 25.4% versus 10.3%) and, to a lesser extent, the nuclear or perinuclear categories (Patterns 1 and 2; 19.5% versus 16.1%). The cell-type composition of each pattern category differed (Figure 5c): cells in the peripheral category were predominantly tumor epithelial cells (about 73%), whereas tumor epithelial cells made up 29-47% of cells in the other categories, in which fibroblasts and other stromal cells were more common. The higher proportion of peripheral components in Xenium may therefore reflect localization patterns specific to tumor cells, while the nuclear and perinuclear components arose largely from stromal cells.

#### Examples of localization subpopulations

Figures 7a and 7b illustrate distinct radial localization patterns identified by ELLA-mix for two genes. *CNN1*, encoding the actinbinding protein calponin 1, exhibited three localization patterns across 284 cells 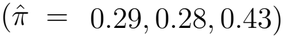, ranging from perinuclear to boundary-enriched distributions (Figure 7a). Given the role of CNN1 in cytoskeletal organization and ovarian cancer progression, this heterogeneity may reflect differences in cytoskeletal organization across cells (Wang et al., 2017). *CCN1*, which encodes a secreted matricellular protein involved in cell-matrix interactions, exhibited two localization patterns across 1,068 cells 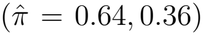, with transcripts concentrated in the mid-cytoplasm or shifted toward the cell boundary (Figure 7b). These differences may be associated with the spatial organization of the secretory pathway, consistent with the localization of mRNAs encoding secreted proteins to the endoplasmic reticulum (Lau, 2011; Lerner et al., 2003). Example cells from each subpopulation are shown in Figure 7c,d. To assess whether the subpopulations were distinct groups of cells, we compared their cell-type composition (Figure 7e). For both genes, all subpopulations came fromresults the same cell type: smooth muscle cells or pericytes for *CNN1* and fibroblasts for *CCN1*. Differential expression analysis between subpopulations identified no significantly differentially expressed genes for either gene. Thus, these subpopulations differed in mRNA localization rather than in cell type or expression program.

**Figure 7.**
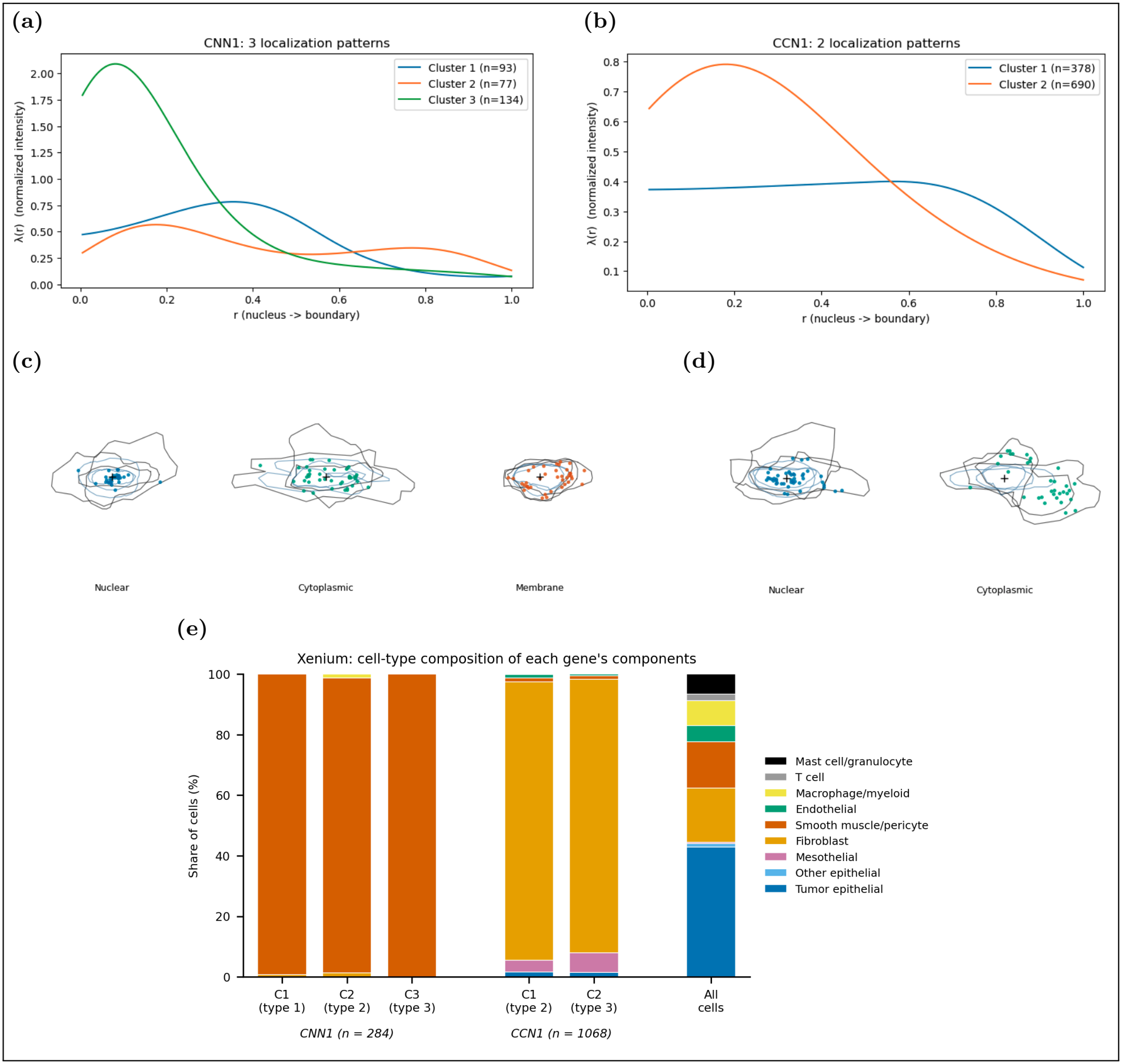
(a) *CNN1* (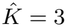; 284 cells). (b) *CCN1* (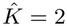; 1,068 cells). (c, d) Example cells from each ELLA-mix subpopulation for *CNN1* (c) and *CCN1* (d). Each panel shows one cell with its boundary, nucleus and the gene’s transcripts, coloured by subpopulation. (e) Cell-type composition of the ELLA-mix subpopulations (C1–C3, with pattern type in parentheses) of *CNN1* and *CCN1*, compared with all cells in the Xenium data. *n* is the number of cells analysed for each gene.

#### Peripheral mRNA localization is associated with proliferative cell states

We investigated the association between radial mRNA localization and cell proliferation using three indicators: detection of *MKI67* or *TOP2A* transcripts, and membership in expression clusters annotated as S phase or G2/M phase (Supplementary Section S3). For each of the 63 genes with multiple localization patterns, we compared the proportions of indicator-positive cells between its outermost and innermost localization subpopulations, using a two-sided sign test to assess whether the direction of the difference was consistent across genes. The outermost subpopulation had a higher proportion of proliferating cells in 39 of 57 testable genes (*p* = 0.008). A similar pattern was observed for S-phase cells (23 of 30 genes; *p* = 0.005), whereas G2/M cells showed no consistent direction (15 versus 14 genes; *p* = 1.0). Gene-level association tests provided additional evidence of proliferation-associated localization heterogeneity for *H3F3B* and *ADAMTS1* (Supplementary Section S3). Previous studies have linked mRNA compartmentalization to cellular quiescence and peripheral mRNA translation to cancer-cell proliferation (Rossi et al., 2026; Dermit et al., 2020), providing a possible biological context for these findings. However, differences in cell-type composition may also contribute to the observed associations. For *H3F3B* and *ADAMTS1*, the outermost (peripheral) subpopulation consisted almost entirely of tumor epithelial cells (94% and 99.6%), whereas the innermost subpopulation was mainly stromal, with tumor epithelial cells making up about 29% and 23% (Figure 8c). For these genes, the association between peripheral localization and proliferation therefore at least partly reflects differences in cell type.

**Figure 8.**
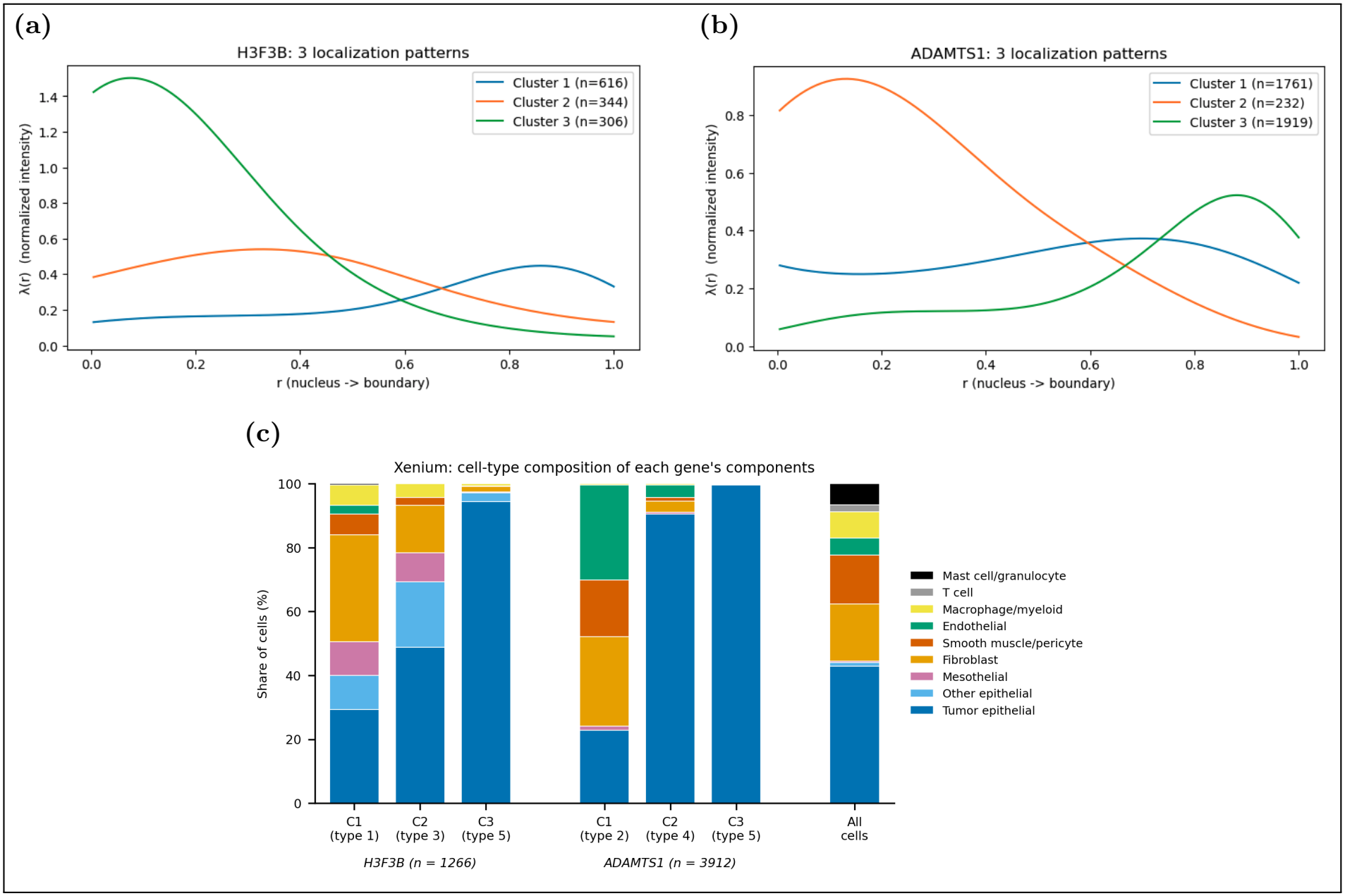
(a) *H3F3B* (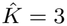; 1266 cells). (b) *ADAMTS1* (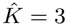; 3912 cells). (c) Cell-type composition of the ELLA-mix subpopulations (C1–C3, with pattern type in parentheses) of *H3F3B* and *ADAMTS1*, compared with all cells in the Xenium data. *n* is the number of cells analysed for each gene.

## 6. Discussion

We developed a mixture model to detect cell-to-cell heterogeneity in subcellular mRNA localization. Building on ELLA (Wang and Zhou, 2025), which estimates a shared radial localization pattern across cells, the proposed model allows different groups of cells to follow distinct patterns without requiring predefined group labels. The model uses flexible spline-based densities to characterize each pattern and estimates the corresponding cell memberships through an EM algorithm. By conditioning on the observed transcript counts, it focuses on differences in spatial distribution rather than expression abundance. An important distinction is that a gene may exhibit nonuniform localization without showing heterogeneity across cells. Our approach addresses this distinction by testing whether multiple localization patterns are supported beyond a flexible single-pattern model.

We developed a model selection procedure that chooses the number of patterns by BIC, considering only models whose clusters are stable across initializations and whose component densities are well separated. Simulations showed that the procedure rarely split single-pattern genes and accurately recovered the number and shapes of localization patterns under the settings considered. Applications to two spatial transcriptomics datasets further demon-strated its utility. In seqFISH+ fibroblasts, we identified distinct localization subpopulations for genes previously characterized by a single localization class. Functional enrichment analysis showed that genes involved in the secretory pathway were more likely to exhibit localization heterogeneity. In Xenium ovarian cancer tissue, we found that subpopulations with transcripts distributed closer to the cell boundary tended to contain higher proportions of proliferating cells, particularly those associated with S phase. These results illustrate how accounting for variation across cells can reveal biological associations that may be obscured when localization is summarized by a single pattern per gene.

The proposed method has several limitations. First, it considers only radial localization and cannot distinguish patterns with different angular distributions. Second, the finite mixture model represents cell-to-cell variation through discrete subpopulations, although localization may vary continuously across cells. We currently consider up to three components, and the stability and separation criteria may limit the detection of smaller or less distinct subpopulations. The analysis also depends on accurate cell segmentation and nucleus localization, and genes with low transcript counts provide limited information for distinguishing patterns. In addition, the simulations revealed a small estimation bias near the nucleus, where relatively few transcripts are available to estimate the radial density. Finally, the associations with proliferation observed in the Xenium data may be partly explained by differences in cell-type composition and should not be interpreted as evidence of a direct relationship.

Several extensions are worth exploring. Incorporating angular information would allow the model to capture directional localization patterns that cannot be represented by radial coordinates alone. Cell-level covariates, including cell type and cell-cycle state, could also be incorporated into the mixture proportions to investigate factors associated with localization heterogeneity. Other extensions include accounting for uncertainty in cell segmentation and transcript positions, and developing more flexible models for continuous variation in localization patterns. These directions would allow a more detailed investigation of how subcellular RNA localization varies across cells and biological conditions.

## References

10x Genomics (2024). FFPE Human Ovarian Cancer with 5K Human Pan Tissue and Pathways Panel plus 100 Custom Genes. Xenium In Situ Gene Expression dataset. Published December 17, 2024.

10x Genomics (2026a). Visium HD spatial gene expression. Available at: https://www.10xgenomics.com/platforms/visium (accessed September 11, 2026).

10x Genomics (2026b). Xenium in situ. Available at: https://www.10xgenomics.com/platforms/xenium (accessed September 11, 2026).

Bierman, R., Dave, J. M., Greif, D. M., and Salzman, J. (2024). Statistical analysis supports pervasive RNA subcellular localization and alternative 3’ UTR regulation. eLife 12, RP87517.

Chen, A. et al. (2022). Spatiotemporal transcriptomic atlas of mouse organogenesis using DNA nanoball-patterned arrays. Cell 185, 1777–1792.e21.

Chen, K. H., Boettiger, A. N., Moffitt, J. R., Wang, S., and Zhuang, X. (2015). Spatially resolved, highly multiplexed RNA profiling in single cells. Science 348, aaa6090.

Das, S., Vera, M., Gandin, V., Singer, R. H., and Tutucci, E. (2021). Intracellular mrna transport and localized translation. Nature reviews Molecular cell biology 22, 483–504.

Dermit, M., Dodel, M., Lee, F. C. Y., Azman, M. S., Schwenzer, H., Jones, J. L., Blagden, S. P., Ule, J., and Mardakheh, F. K. (2020). Subcellular mrna localization regulates ribosome biogenesis in migrating cells. Developmental Cell 55, 298–313.e10.

Eng, C.-H. L., Lawson, M., Zhu, Q., Dries, R., Koulena, N., Takei, Y., Yun, J., Cronin, C., Karp, C., Yuan, G.-C., and Cai, L. (2019). Transcriptome-scale super-resolved imaging in tissues by RNA seqFISH+. Nature 568, 235–239.

Lau, L. F. (2011). CCN1/CYR61: The very model of a modern matricellular protein. Cellular and Molecular Life Sciences 68, 3149–3163.

Lerner, R. S., Seiser, R. M., Zheng, T., Lager, P. J., Reedy, M. C., Keene, J. D., and Nicchitta, C. V. (2003). Partitioning and translation of mRNAs encoding soluble proteins on membrane-bound ribosomes. RNA 9, 1123–1137.

Mah, C. K., Ahmed, N., Lopez, N. A., Lam, D. C., Pong, A., Monell, A., Kern, C., Han, Y., Prasad, G., Cesnik, A. J., Lundberg, E., Zhu, Q., Carter, H., and Yeo, G. W. (2024). Bento: a toolkit for subcellular analysis of spatial transcriptomics data. Genome Biology 25, 82.

NanoString Technologies (2026). CosMx spatial molecular imager. Available at: https://nanostring.com/products/cosmx-spatial-molecular-imager/ (accessed September 11, 2026).

Novoselsky, R., Golani, O., Barkai, T., Kedmi, M., Goliand, I., Fine, M., Kent, I., Nachmany, I., and Itzkovitz, S. (2026). Subcellular mrna localization patterns across tissues resolved with spatial transcriptomics. Nature Communications 17, 5466.

Rossi, A., Coum, A., Madelenat, M., Harris, L., Strohbuecker, S., Chai, A., Fiaz, H., Chaouni, R., Faull, P., Heaven, N. C., Grey, W., Bonnet, D., Hamid, F., Makeyev, E. V., Snijders, A. P., Kelly, G., Guillemot, F., and Sousa-Nunes, R. (2026). Cellular quiescence uncouples the proteome from the transcriptome in neural stem cells. The EMBO Journal 45, 1697–1727.

Shestakova, E. A., Singer, R. H., and Condeelis, J. (2001). The physiological significance of β-actin mRNA localization in determining cell polarity and directional motility. Proceedings of the National Academy of Sciences of the United States of America 98, 7045–7050.

Walter, F. C., Stegle, O., and Velten, B. (2023). FISHFactor: a probabilistic factor model for spatial transcriptomics data with subcellular resolution. Bioinformatics 39, btad183.

Wan, H., Yang, P., Hou, S., Wang, J., and Zhou, X. (2026). Svc: A vision transformer-based spatially and virtually embedded cell model for deciphering subcellular spatial transcriptomic heterogeneity. bioRxiv pages 2026–10.

Wang, J. X. and Zhou, X. (2025). ELLA: modeling subcellular spatial variation of gene expression within cells in high-resolution spatial transcriptomics. Nature Communications 16, 9920.

Wang, K.-H., Chu, S.-C., and Chu, T.-Y. (2017). Loss of calponin h1 confers anoikis resistance and tumor progression in the development of high-grade serous carcinoma originating from the fallopian tube epithelium. Oncotarget 8, 61133–61145.

Wouters, D. et al. (2026). SubCellSpace: Automated characterization of subcellular mRNA localization patterns in spatial transcriptomics. bioRxiv Preprint.

Zhao, Y., Shapiro, S. S., and Eto, M. (2016). F-actin clustering and cell dysmotility induced by the pathological w148r missense mutation of filamin b at the actin-binding domain. American Journal of Physiology-Cell Physiology 310, C89–C98.

