## Supplementary Material for "ELLA-mix: Detecting Cell-to-Cell Heterogeneity in Subcellular RNA Localization"

### Supplementary Materials for "ELLA-mix: Detecting Cell-to-Cell Heterogeneity in Subcellular RNA Localization"

#### S1 *Additional details for model selection*

*Gene filtering and implementation.* We restrict the analysis to genes detected in at least 20 cells with at least 10 transcripts each. For each eligible gene, we consider mixture models with up to $K = 3$ components, using $P = 5$ cubic B-spline basis functions with an interior knot at 0.5. Each candidate model is fitted from three random initializations, and the fit with the highest observed-data log-likelihood is retained. The cell assignments from all three initializations are used to assess clustering stability.

*Clustering stability.* For a candidate model with $k \geqslant 2$ components, we assess the stability of cell assignments across random initializations using the adjusted Rand index (ARI). Let $\widehat{\mathbf{z}}_k^{(r)}$ denote the vector of cell assignments obtained from initialization $r$, where each cell is assigned to the component with the highest posterior membership probability. The minimum pairwise ARI is defined as

$$\mathrm{ARI}_{\min}(k) = \min_{1 \leqslant r < s \leqslant 3} \mathrm{ARI}\left(\widehat{\mathbf{z}}_k^{(r)}, \widehat{\mathbf{z}}_k^{(s)}\right). \tag{S1}$$

An ARI of 1 indicates identical partitions, whereas values near 0 indicate agreement no better than chance. We require $\mathrm{ARI}_{\min}(k) > 0.80$ for a model to satisfy the stability criterion.

*Component separation.* To assess whether the fitted components represent distinct localization patterns, we use the total variation (TV) distance between their radial densities. For components $a$ and $b$, the TV distance is

$$\mathrm{TV}(a, b) = \frac{1}{2} \int_0^1 \left| f_a(\rho; \widehat{\boldsymbol{\beta}}_a) - f_b(\rho; \widehat{\boldsymbol{\beta}}_b) \right| \, d\mu(\rho). \tag{S2}$$

We evaluate this distance numerically on a grid of $G = 200$ equally spaced radial points $r_1, \ldots, r_G$ in $[0, 1]$. Let $w_g = 2\pi r_g \Delta r$ denote the radial area weight at grid point $r_g$. The corresponding normalized probability mass for component $a$ is

$$p_a(r_g) = \frac{w_g f_a(r_g; \widehat{\boldsymbol{\beta}}_a)}{\sum_{h=1}^{G} w_h f_a(r_h; \widehat{\boldsymbol{\beta}}_a)}. \tag{S3}$$

The TV distance is approximated by

$$\mathrm{TV}(a, b) \approx \frac{1}{2} \sum_{g=1}^{G} |p_a(r_g) - p_b(r_g)|. \tag{S4}$$

For a model with $k \geqslant 2$ components, we require the minimum pairwise TV distance to be at least 0.20. This corresponds to an overlap of at most 80% between any two fitted component densities.

*Admissibility and model selection.* A candidate model with $k \geqslant 2$ is admissible if both the stability and separation criteria are satisfied:

$$A_k = I\left\{\mathrm{ARI}_{\min}(k) > 0.80 \text{ and } \min_{a<b} \mathrm{TV}(a, b) \geqslant 0.20\right\}. \tag{S5}$$

We set $A_1 = 1$. Among the admissible models, the number of components is selected by minimizing the BIC defined in Section 3.3 of the main text, $\mathrm{BIC}_k = -2\hat{\ell}_k + p_k \log n$. With $P = 5$ basis functions, the number of identifiable parameters is $p_k = 5k - 1$, giving 4, 9, and 14 parameters for $k = 1, 2$, and 3, respectively.

#### S2 *Additional simulation details*

*Data generation.* We generated synthetic genes with known radial localization patterns, representing each cell by the unit disk. For the component-selection simulations, we simulated 150 cells per gene in the main setting and 50, 100 and 200 cells per gene as additional settings. Transcript counts per cell were drawn from a negative binomial distribution with mean 6 and size 2, and, as in the real data analysis, only cells with at least 10 transcripts were retained. Retained cells contained an average of 13.9 transcripts, close to the median of 14 in the seqFISH+ dataset.

Under complete spatial randomness (CSR), transcripts are uniform over the unit disk, with radial density $g_0(r) = 2r$. Localized patterns mixed CSR with a radial bump,

$$g_{p,s}(r) = (1-s)\,2r + s\,h_p(r), \qquad h_p(r) \propto r\exp\left\{-\frac{(r-\mu_p)^2}{2\sigma_p^2}\right\}, \qquad 0 \leqslant r \leqslant 1, \qquad \text{(S6)}$$

where $\mu_p$ and $\sigma_p$ set the location and spread of pattern $p$, and $s$ is the localization strength. The factor $r$ in $h_p$ keeps the spatial intensity $h_p(r)/(2\pi r)$ finite at the cell center. We used three patterns: nuclear ($\mu = 0.15$), cytoplasmic ($\mu = 0.50$), and membrane ($\mu = 0.80$), each with $\sigma = 0.173$. Unless otherwise stated, $s = 0.472$, giving TV distances from CSR of 0.32, 0.17 and 0.13, respectively, and pairwise TV distances of 0.30 (nuclear–cytoplasmic), 0.27 (cytoplasmic–membrane) and 0.43 (nuclear–membrane). The closest pair is thus comparable to the average separation between the closest fitted components of three-pattern genes in the real data (0.25). All cells sharing a pattern followed the same radial density.

*Testing for nonuniform localization.* Let $\hat{\ell}_{\mathrm{CSR}}$ and $\hat{\ell}_1$ denote the maximized conditional log-likelihoods under CSR and the single-component spline model. The likelihood-ratio statistic $T = 2(\hat{\ell}_1 - \hat{\ell}_{\mathrm{CSR}})$ was compared with its asymptotic $\chi^2_{P-1}$ null distribution, that is, $\chi^2_4$ for $P = 5$ cubic B-spline basis functions.

For type I error, we generated 300 CSR genes for each of 50, 100, 150 and 200 cells per gene and recorded the proportion rejected at level 0.05, along with the distribution of $p$-values. For power, we varied $s \in \{0.05, 0.10, 0.15, 0.20, 0.30, 0.40, 0.60, 1\}$ in (S6) for each pattern, giving TV distances from CSR between 0.013 and 0.68, and simulated 200 genes per pattern, strength and number of cells, together with 200 CSR genes per number of cells. Power was the proportion of genes rejected at level 0.05.

*Selection of the number of patterns.* We considered eight scenarios with 300 genes each: CSR, the three single patterns, the three pairs (nuclear–cytoplasmic, nuclear–membrane, cytoplasmic–membrane) and all three patterns. Two-pattern scenarios assigned 65% and 35% of cells to the first and second listed patterns; the three-pattern scenario assigned 24%,

47% and 29% to the nuclear, cytoplasmic and membrane patterns. For each gene, we fitted models with one to three components, using three random initializations each, and selected the number of components by minimizing BIC among admissible models (Section 3.3 of the main text and Section S1). Selection accuracy was the proportion of genes for which the selected number equaled the true number.

*Accuracy of estimated localization patterns.* For genes with the correctly selected number of components $K$, we compared each fitted radial density $\widehat{g}_k$ with the true generating density $g_k$ using the TV distance $\mathrm{TV}(g,h) = \frac{1}{2}\int_0^1 |g(r) - h(r)|\,dr$. Because component labels are arbitrary, fitted components were matched to true patterns by the permutation $\sigma$ minimizing the total TV distance, and estimation error was the average over matched pairs,

$$E_{\mathrm{TV}} = \min_{\sigma} \frac{1}{K} \sum_{k=1}^{K} \mathrm{TV}\big(\widehat{g}_{\sigma(k)}, g_k\big). \tag{S7}$$

We summarized $E_{\mathrm{TV}}$ by scenario and number of cells per gene.

S3 *Assessment of cell proliferation*

We analysed the 63 genes with multiple localization patterns ($\hat{K} = 2$ for 48 genes and $\hat{K} = 3$ for 15). Across the 147,366 cells in which at least one of these genes was analysed, 23.6% were marker-positive (*MKI67*, 9.1%; *TOP2A*, 19.8%), 13.6% belonged to the S-phase cluster, and 8.6% to the G2/M cluster. Because *TOP2A* was itself one of the analysed genes, its marker indicator was based on *MKI67* alone.

For each gene and indicator, we calculated the proportion of indicator-positive cells within each localization subpopulation. A gene was considered testable for an indicator if it had at least five indicator-positive cells; this gave 57 testable genes for the marker indicator and 30 each for the S-phase and G2/M indicators. To assess directional consistency across genes, we compared the proportions between the outermost and innermost subpopulations, defined by the mean radial positions of their fitted localization patterns, and recorded the sign of the difference. We then applied a two-sided exact sign test to assess whether positive and negative differences occurred with equal frequency across testable genes; genes with equal proportions (one gene, for G2/M) were excluded from the sign test.

As a complementary analysis, we tested the association between localization subpopulation membership and each proliferation indicator for each testable gene, using Fisher's exact test for genes with two subpopulations and the chi-squared test for genes with three, with a permutation $p$-value when expected counts were small. We adjusted the resulting gene-level $p$-values for multiple testing using the Benjamini–Hochberg procedure; 10, 14 and 9 genes had $q < 0.05$ for the marker, S-phase and G2/M indicators, respectively.
